# The demographic history and mutational load of African hunter-gatherers and farmers

**DOI:** 10.1101/131219

**Authors:** Marie Lopez, Athanasios Kousathanas, Hélène Quach, Christine Harmant, Patrick Mouguiama-Daouda, Jean-Marie Hombert, Alain Froment, George H. Perry, Luis B. Barreiro, Paul Verdu, Etienne Patin, Lluís Quintana-Murci

## Abstract

The distribution of deleterious genetic variation across human populations is a key issue in evolutionary biology and medical genetics. However, the impact of different modes of subsistence on recent changes in population size, patterns of gene flow, and deleterious mutational load remains unclear. Here, we report high-coverage exome sequencing data from various populations of rainforest hunter-gatherers and farmers from central Africa. We find that the recent demographic histories of hunter-gatherers and farmers differed considerably, with population collapses for hunter-gatherers and expansions for farmers, accompanied by increased gene flow. We show that purifying selection against deleterious alleles is of similar efficiency across African populations, in contrast with Europeans where we detect weaker purifying selection. Furthermore, the per-individual mutation load of rainforest hunter-gatherers is similar to that of farmers, under both additive and recessive models. Our results indicate that differences in the cultural practices and demographic regimes of African populations have not resulted in large differences in mutational burden, and highlight the beneficial role of gene flow in reshaping the distribution of deleterious genetic variation across human populations.

## Introduction

Human populations have undergone radical changes in size over the last 100,000 years, due to various range expansions, bottlenecks, and periods of rapid growth^1-3^. An understanding of the ways in which these demographic changes have affected the ability of human populations to purge deleterious variants is crucial for the dissection of the genetic architecture of human diseases^4-8^. Theoretical population genetic studies have shown that most new mutations resulting in amino acid substitutions are rapidly culled from populations through purifying selection, at a rate dependent on both the effective population size (*N*_*e*_) and the distribution of selection coefficients (*s*)^9^. Indeed, new mutations with deleterious effects are less efficiently purged from populations with a small *N*_*e*_, in which genetic drift has a stronger effect than selection.

Recent empirical studies based on population genomic-scale datasets have revealed an unsuspectedly large burden of rare and low-frequency amino acid-altering variants in the human genome^10-14^. Population genetic studies have also reported differences in the number, frequency and distribution of putatively deleterious variants across populations, and it has been suggested that these differences result from demographic events, including explosive growth, founder events, bottlenecks, and inbreeding^8,15-22^. For example, the higher proportion of deleterious variants detected in non-Africans has been interpreted as the result of a decrease in selection efficacy after the out-of-Africa bottleneck and recent explosive growth^15,22^. It has also been suggested that the mutation load, *i.e.*, the difference between the theoretical optimal fitness and actual fitness of a population, increases with distance from Africa due to serial founder effects^17^.

By contrast, other studies have reported no detectable differences between populations for summary statistics approximating the per-individual mutation load, consistent with an equal efficiency of purifying selection across populations, at least for an additive dominance (semidominant) model^23,24^. Several factors underlie these apparently conflicting results, including differences in the statistics used to evaluate selection efficacy and to approximate mutation load, the methods used to assess the significance of population differences, and the choice of predictive algorithms for defining deleteriousness^25-28^. Most studies have compared statistics across populations without integrating explicit models of demography and selection, despite the availability of methods for estimating the distribution of fitness effects of new nonsynonymous mutations (DFE) in a population, accounting for its demographic history^29,30^.

In addition to these methodological considerations, it remains largely unknown how differences in cultural practices between populations have affected their demographic regimes, and the efficiency of purifying selection. About 5% of human populations currently rely on modes of subsistence not associated with recent population growth, such as hunting and gathering^31^. Africa has the largest group of hunter-gatherer populations, the rainforest hunter-gatherers (known as “pygmies”), who have traditionally lived in small, mobile groups scattered across the central African equatorial forest^32^. By contrast, the neighboring sedentary populations practice agriculture, and are known to have recently expanded across sub-Saharan Africa^33^. These contrasting lifestyles are associated with differences in the demographic histories of these populations, but some aspects of the genetic and demographic histories of rainforest hunter-gatherers and farmers remain unclear, given the highly heterogeneous nature of the genetic data used and the limited sample sizes included in earlier studies^34-39^. Furthermore, signatures of recent demographic change can be detected only if the full site frequency spectrum is obtained from sequencing data for relatively large sample sizes, a condition not met by any of the previous studies of these populations.

Here we aimed to understand how differences in demographic history might have affected the efficacy of purifying selection and the corresponding mutational load in populations with different subsistence strategies. We generated 300 high-coverage exomes from various rainforest hunter-gatherer and farming populations from both western and eastern central Africa, a dataset that was supplemented with 100 exomes from a European population. Using a coalescent-based composite likelihood approach, we first estimated the demographic parameters characterizing these populations in terms of changes in population size, split times and gene flow. We then compared the efficiency of selection to purge deleterious alleles and the estimated deleterious mutational load across populations, with the aim of improving our understanding of the effects of traditional lifestyle and demographic history on the mutational burden of contemporary human populations.

## Results

### Population Exome Sequencing Dataset

We constituted a unique collection of central African populations with different historical lifestyles: 100 Baka mobile rainforest hunter-gatherers (wRHG) and 100 Nzebi and Bapunu sedentary farmers (wAGR) from western central Africa (*i.e,.* Gabon and Cameroon), and 50 BaTwa rainforest hunter-gatherers (eRHG) and 50 BaKiga farmers (eAGR) from eastern central Africa (*i.e.,* Uganda) (Fig. 1a and Supplementary Table 1). We first investigated the genetic structure, and potential substructure, of the study populations, using genome-wide SNP data for the 300 individuals (Supplementary Note). ADMIXTURE^40^ and principal component analysis (PCA)^41^ separated African populations on the basis of mode of subsistence (AGR *vs.* RHG), before splitting RHG into western and eastern groups (Fig. 1b and Supplementary Figs 1 and 2)^34,36,37^. The inbreeding coefficient (*F*_IS_) distributions and additional ADMIXTURE analyses provided no evidence of internal substructure within the groups studied (Supplementary Figs 3 and 4).

**Figure 1.**
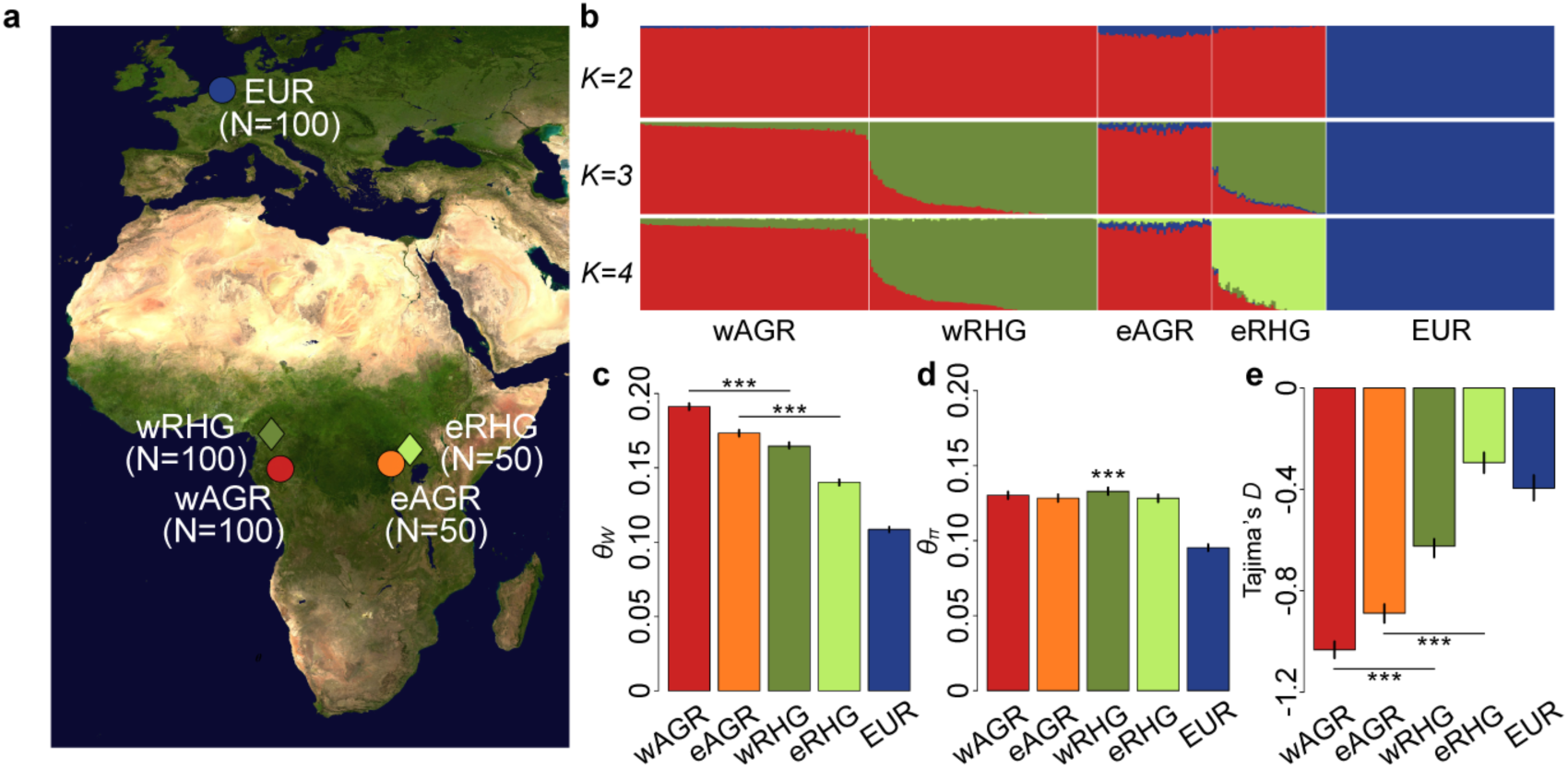
Genetic structure and diversity of African rainforest hunter-gatherers and farmers. The studied populations were African rainforest hunter-gatherers (RHG), neighboring agriculturalists (AGR) and Europeans (EUR). **a**, Location of the sampled populations; **b**, Estimation of ancestry proportions with the clustering algorithm ADMIXTURE using the SNP array data; **c**, Watterson’s estimator *θ*_*W*_; **d**, Pairwise nucleotide diversity *θ*_*π*_; **e**, Tajima’s *D*. **c**, **d**, **e,** all neutrality statistics were calculated with exome sequencing data for 4-fold degenerate synonymous sites and confidence intervals were obtained by bootstrapping by site. Significance was assessed between wAGR/wRHG and eAGR/eRHG in **c** and **e**, and for all comparisons in **d**. ***: *P*-value < 10^−3^.

We then performed whole-exome sequencing for the entire collection of 300 unrelated individuals at high coverage (mean depth 68x). We identified 406,270 quality-filtered variants, including 67,037 newly identified variants (Supplementary Table 1 and Supplementary Figs 5 and 6). For calibration of the demographic inferences for African populations, and comparison with a well-studied population for mutation load^15,23,24^, we supplemented our dataset with high-coverage exome sequences from 100 Belgians of European ancestry (EUR) generated with the same experimental and analytical procedures^42^, yielding a final dataset of 488,653 SNPs.

We first sought to obtain a broad view of diversity in RHG and AGR populations, by calculating neutral diversity statistics for synonymous variants (4-fold degenerate variants). Watterson’s *θ* (*θ*_*W*_) was significantly higher in the western and eastern AGR populations than in the RHG populations (*P*-value < 10^−3^; Fig. 1c and Supplementary Table 2), due to the larger proportion of low-frequency variants, as demonstrated by the significantly more negative Tajima’s *D* value obtained (*P*-value < 10^−3^ for both comparisons; Fig. 1e). However, western RHG had the highest pairwise nucleotide diversity *θ*_*π*_ (*P*-value < 10^−3^ for all comparisons; Fig. 1d), suggesting a large historical *N*_*e*_ for this population. These results indicate that the African populations studied have similar levels of genetic diversity, regardless of their historical mode of subsistence, but their different allele frequency distributions suggest contrasting demographic histories.

### Model-based Inference of Population Divergence Times, Size Changes and Gene Flow

We then evaluated how differences in subsistence strategies between RHG and AGR populations have affected their demographic histories. Demographic parameters were estimated by fitting models incorporating all the populations studied, including EUR, into pairwise (2D) site frequency spectra (SFS), with the coalescent-based composite likelihood approach *fastsimcoal2*^43^ (Supplementary Note). Going forward in time, we assumed an early population size change for the ancestor of all populations, followed by size changes coinciding with population splits, and an additional population size change for EUR (Supplementary Fig. 7). Because the chronology of divergences between these populations remains unknown, we formulated three branching models, each assuming that a different population (EUR, RHG or AGR) was the first to split off from the remaining groups (*i.e.*, EUR-first, RHG-first, AGR-first, Fig. 2). Furthermore, admixture between wRHG and wAGR, eRHG and eAGR, and between eAGR and EUR has been documented (Fig. 1b and refs.^39,44,45^). We therefore estimated parameters by considering two epochs of continuous migration between population pairs, allowing for asymmetric gene flow.

**Figure 2.**
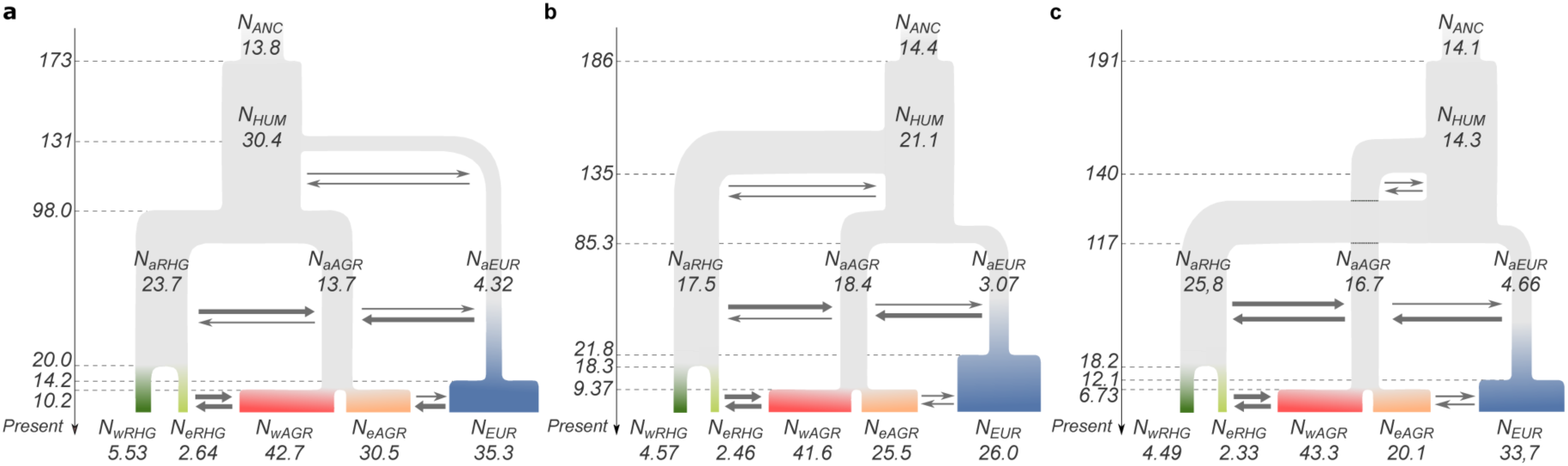
Inferred demographic models of the studied populations. **a**, EUR-first branching model, in which the European population diverged from African populations before the divergence of the ancestors of RHG (aRHG) and AGR (aAGR); **b**, RHG-first branching model, in which the aRHG was the first to diverge from the other groups; and **c**, AGR-first branching model, in which aAGR was the first to diverge from the other groups. We assumed an ancient change in the size of the ancestral population of all humans (ANC). We assumed that each subsequent divergence of populations was followed by an instantaneous change in effective population size (*N*_*e*_). We also assumed that there were two epochs of migration between the following population pairs: wAGR/aAGR and wRHG/aRHG, eAGR/aAGR and eRHG/aRHG, and EUR and eAGR/aAGR. The figure labels correspond to the estimated parameters of the model (Supplementary Table 4). Bold arrows indicate 2*Nm* > 1.

The three branching models produced non-significant differences in likelihood (*P*-value > 0.05 for all comparisons; Supplementary Table 3), and all three models fitted both observed marginal 1D SFS and *F*_ST_ values well (Supplementary Figs 8 and 9). These models consistently provided similar estimates for key demographic parameters (Supplementary Table 4). Our results suggest that the ancestors of the contemporary RHG, AGR and EUR populations diverged between 85 and 140 thousand years ago (kya), from an ancestral population that underwent demographic expansion between 173 and 191 kya (*T*_*ANC*_) (Fig. 2). After the initial population splits, the *N*_*e*_ of AGR and RHG (*N*_*aAGR*_ and *N*_*aRHG*_) remained within a range extending from 0.55 to 2.2 times the ancestral African *N*_*e*_ (*N*_*HUM*_), whereas EUR (*N*_*aEUR*_) experienced a decrease in *N*_e_ by a factor of three to seven (Supplementary Tables 4 and 5). The ancestors of the wRHG and eRHG populations diverged 18 to 20 kya (*T*_*RHG*_), and underwent a decreased in *N*_e_ by a factor of 3.8 to 5.7 for the wRHG (*N*_*wRHG*_) and 7.1 to 11 for the eRHG (*N*_*eRHG*_), regardless of the branching model considered. The ancestors of the AGR (*N*_*aAGR*_) split into western and eastern populations 6.7 to 11 kya (*T*_*AGR*_), and underwent a mild expansion, by a factor of 2.3 to 3.1 for the wAGR (*N*_*wAGR*_) and 1.2 to 2.2 for the eAGR (*N*_*eAGR*_). The EUR population experienced a 7.1- to 8.3-fold expansion (*N*_*EUR*_) 12 to 22 kya (*T*_*EUR*_).

The 95% confidence intervals of estimated parameters were generally wide (Supplementary Table 4), owing to the complexity of the models and the limited number of 4-fold degenerate synonymous sites used. Nevertheless, we obtained significant support, based on ratios of ancestral to current *N*_e_, for a recent bottleneck in both wRHG and eRHG populations, and for a recent expansion of wAGR and EUR populations, for all branching models (*i.e.*, *N*_*aRHG*_/*N*_*RHG*_ > 1, and *N*_*aAGR*_/*N*_*wAGR*_ and *N*_*aEUR*_/*N*_*EUR*_ < 1; *P*-value < 0.05; Supplementary Table 5).

The estimated migration parameters were also mostly similar between branching models (Supplementary Table 6). We accounted for differences in *N*_*e*_ between the recipient populations, by comparing the effective strength of migration (2*Nm*) between populations exchanging migrants. For the ancient migration epoch, we found evidence for migration between the ancestors of RHG and AGR (Supplementary Table 4), although we were unable to determine its direction and strength with confidence. During the recent epoch, migration between RHG and AGR was confidently inferred to be strong and mostly symmetric across models (2*Nm* > 17 and 8 in western and eastern groups, respectively). Finally, we inferred that 2*Nm* was larger for migration from EUR to AGR than the opposite direction, for both migration epochs (from 7- to 120-fold stronger across models and epochs; Supplementary Table 4), consistent with back-to-Africa migrations^2,44,45^.

Together, our analyses support a demographic history in which a large ancestral population of RHG continuously exchanged migrants with the ancestors of AGR until about 10,000-20,000 years ago, when the ancestors of the RHG and AGR populations experienced bottlenecks and expansions, respectively, and migration between these two groups increased markedly.

### Comparing the Efficacy of Selection Across Populations

We then explored whether the different inferred demographic histories of RHG and AGR populations affected the efficiency with which deleterious alleles were purged by selection. We used a model-based inference of the distribution of fitness effects (DFE) of new nonsynonymous mutations across populations, using *DFE-α*, which explicitly incorporates models of non-equilibrium demography^46^. We first fitted a three-epoch demographic model to the synonymous SFS per population (Supplementary Table 7), yielding broadly consistent results with those generated by *fastsimcoal2* based on 1D SFS (Supplementary Note and Supplementary Table 8). We then fitted a gamma distribution DFE model to the nonsynonymous SFS, accounting for demography. The fit of the gamma DFE model to the data was good (Spearman’s ρ > 0.8 between the expected and observed SFS; Supplementary Fig. 10).

The inferred parameters indicated an L-shaped DFE with a significantly lower mean, *E*(*N*_*e*_*s*), in Europeans than in Africans, but no significant difference between African populations (Supplementary Table 7). However, it is difficult to estimate the *E*(*N*_*e*_*s*) parameter accurately, given that highly deleterious mutations with a strong impact on this parameter are unlikely to be detected, even in large samples^47^. We therefore also summarized the DFE by computing the proportion of mutations assigned by the inferred gamma distribution into four *N*_*e*_*s* ranges (0-1, 1-10, 10-100 and >100, corresponding to neutral, weakly, moderately and strongly deleterious mutations, respectively), a summary much more accurately estimated by *DFE-α*^46^. We estimated that 38.4% and ∼25% of new mutations were weakly-to-moderately deleterious (*N*_*e*_*s* 1-100) in Europeans and Africans (Fig. 3), respectively, consistent with weaker selection in Europeans^8,15,22,26^.

**Figure 3.**
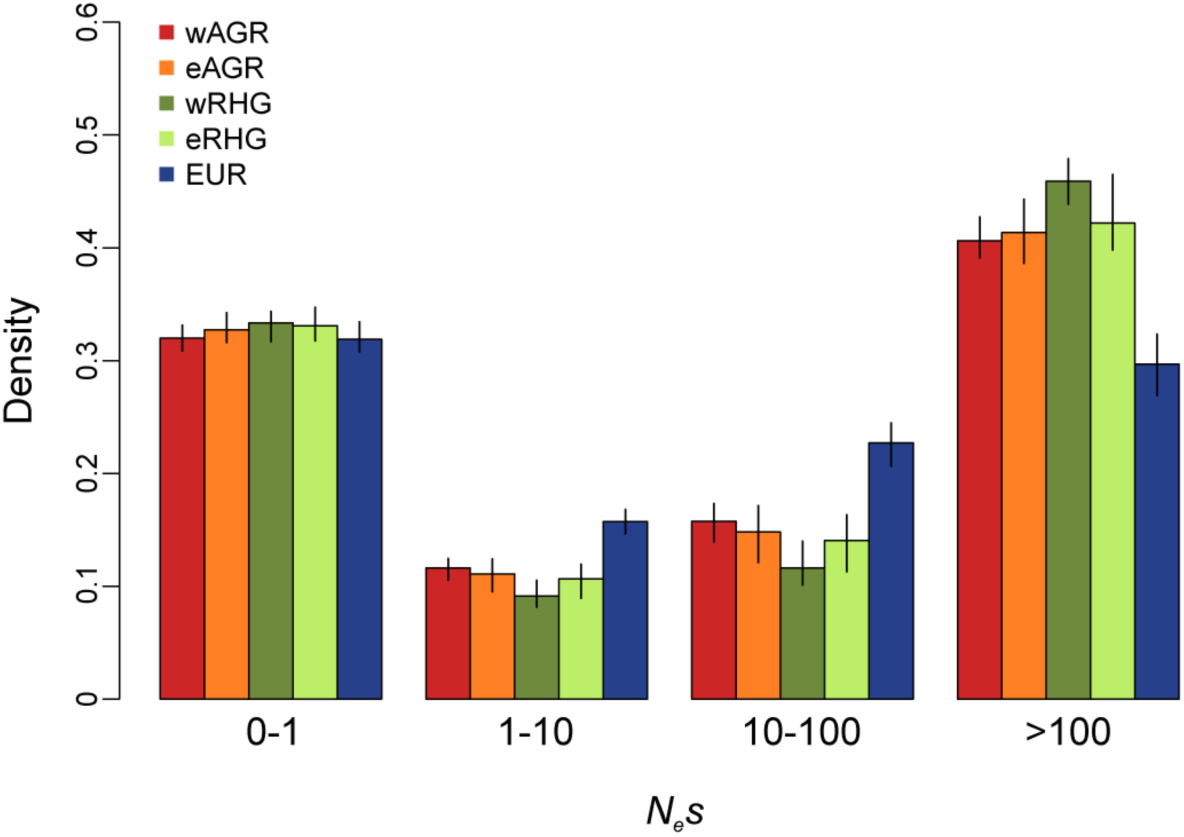
Distribution of fitness effects of new nonsynonymous mutations. The inferred fraction of new mutations in different bins of selection strength (*N*_*e*_*s* = 0-1, 1-10, 10-100, >100) with *DFE-α,* assuming a three-epoch demography fitted for each population separately. We used non-CpG sites and confidence intervals were calculated by bootstrapping by site 100 times.

By contrast, the differences in the density assigned to each *N*_*e*_*s* category were more limited for the RHG and AGR populations, and these differences were not statistically significant (Fig. 3). In particular, we obtained almost identical results with *DoFE*, an alternative method^48^ that does not assume an explicit demographic model and fits nuisance parameters to the synonymous SFS (Supplementary Table 7). Together, our findings indicate that the contrasting demographic regimes of RHG and AGR have not differentially affected the efficacy of selection in these populations.

### Estimating Differences in Mutational Burden Between Populations

The similarity of selection efficacy across African populations suggests that these populations have similar deleterious mutation loads. However, the models used by *DFE-α* assume that deleterious mutations are additive (*i.e.*, semi-dominant) and do not consider individual genotype information, which is crucial for the estimation of recessive mutation load^23,49^. We thus examined the per-individual distribution of heterozygous (*N*_*het*_) and homozygous (*N*_*hom*_) variants and their weighted sum (number of derived alleles, *N*_*alleles*_=*N*_*het*_+2×*N*_*hom*_, refs.^17,23^), these last two variables being monotonically related to the individual load under recessive and additive models of dominance, respectively^23,28^. These statistics were not affected by sequencing quality (Supplementary Fig. 11). The data were partitioned into variants presumed to evolve neutrally (synonymous) and variants likely to be under selective constraints (nonsynonymous variants, variants with Genomic Evolutionary Rate Profiling-Rejected Substitution [GERP RS] scores greater than 4, and loss-of-function mutations), as suggested by their site frequency spectra (Supplementary Fig. 12). We also performed simulations to predict *N*_*hom*_ and *N*_*alleles*_ under neutrality across our three branching models (Supplementary Fig. 13). For both observed genotype counts in all site classes and simulated counts, we used ratios of genotype counts between populations to facilitate comparisons and to make use of the shared history of populations to increase precision.

Population differences in *N*_*alleles*_ values observed in site classes with large numbers of variant sites (synonymous, nonsynonymous and GERP RS >4) and simulated under neutrality did not exceed 2% and were not significantly different for any of the population pairs examined (Fig. 4 and Supplementary Table 9). For LOF mutations, for which there were fewer variant sites, the *N*_*alleles*_ ratio between populations did not exceed 8% and were not significantly different from one. Conversely, *N*_*hom*_ was more than 20% higher in Europeans than in Africans, for both simulated neutral and observed sites in all classes, probably due to the long-term bottleneck experienced by European populations. Much smaller differences in *N*_*hom*_ were observed between RHG and AGR populations; nonsynonymous *N*_*hom*_ was about 2-4% lower in wRHG than in other Africans, and ∼1-4% higher in eRHG than in other African populations, supporting the view that the recent demographic events experienced by these African populations had no major effect on their additive and recessive mutation loads.

**Figure 4.**
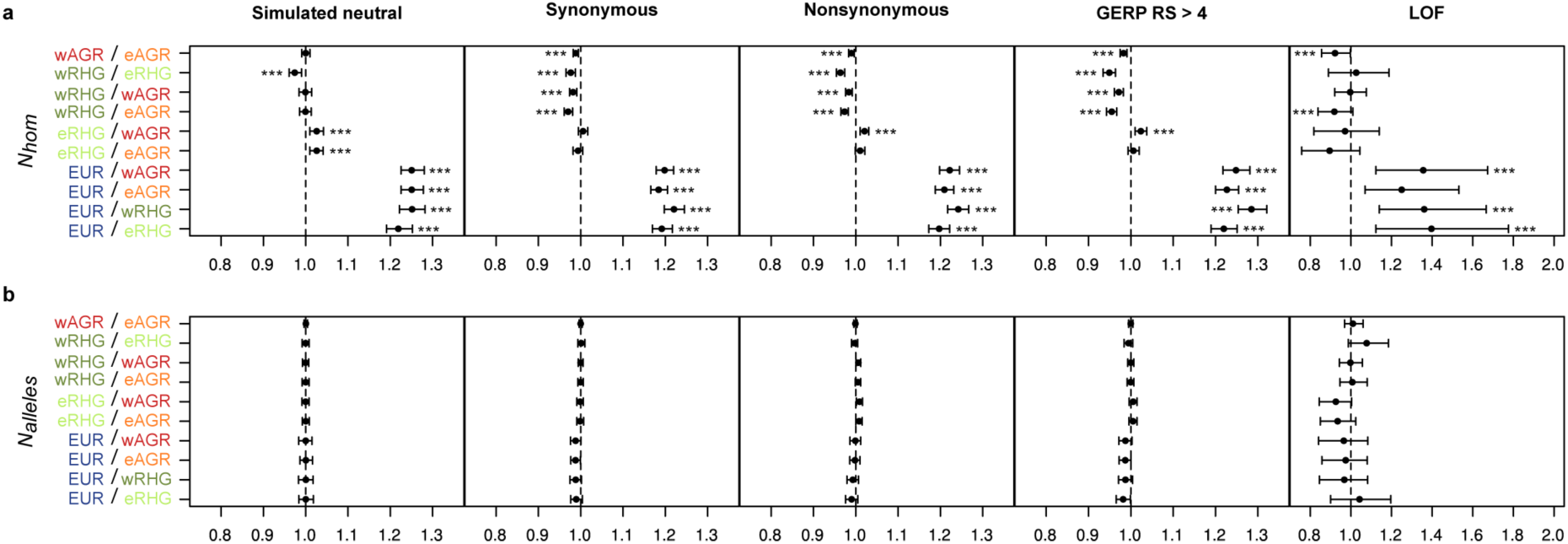
Comparison of approximated recessive and additive mutational loads across human populations. Between-population ratios of the per-individual counts of **a**, homozygous genotypes (*N*_*hom*_) and **b**, numbers of alleles (*N*_*alleles*_=*N*_*het*_+2×*N*_*hom*_). These ratios were obtained through simulations assuming neutrality under the demographic model with the highest likelihood (RHG-first) (panel “Simulated neutral”), and were computed with the observed data for several site classes (panels “Synonymous”, “Nonsynonymous”, “GERP RS >4” and “LOF”). Confidence intervals were calculated by dividing the SNP data into 1000 blocks and carrying out bootstrap resampling of sites 100 times. ***: *P*-value < 10^−3^.

We then investigated the extent to which the similarity of *N*_*hom*_ between RHG and AGR populations could be attributed to the strong gene flow inferred between them. We performed simulations under our best-fitting demographic models, with migration between populations set to zero. The ratio of *N*_*hom*_ between RHG and AGR populations was much higher for simulations without migration (Supplementary Fig. 14) than for simulations with migration (Supplementary Fig. 13), with the strongest effect being observed for eRHG (ratio of *N*_*hom*_ for eRHG relative to eAGR of 1.1 in the absence of migration versus 1.025 with migration). Thus, our simulations suggest that the *N*_*hom*_ for RHG would be higher if there were no gene flow between these populations and AGR populations. Moreover, as RHG individuals have various degrees of AGR ancestry (Fig. 1b), it is possible to test empirically whether the proportion of AGR ancestry is related to proxies of mutation load. We found a significant negative correlation between the nonsynonymous *N*_*hom*_ of eRHG individuals and their estimated AGR ancestry (*P*-value = 4×10^−3^; Supplementary Fig. 15), suggesting that admixture can effectively reduce the recessive mutation load.

### Comparing the Genomic Distribution of Deleterious Variants

Despite the observed similarity between populations in terms of per-individual estimates of mutation load, we explored possible differences between populations in the genomic distribution of putatively deleterious alleles, due to variable levels of inbreeding, leading to extended runs of homozygosity (ROH) (Supplementary Note and Supplementary Fig. 16), or fluctuating selective pressures across gene functional categories. We found that *N*_*hom*_ and *N*_*alleles*_ were higher in ROH than in other regions of the genome, but no significant differences in these patterns were found between populations (Supplementary Fig. 17). However, the lower rates of recombination for ROH (*P*-value < 10^−3^; Supplementary Fig. 18) suggest that the accumulation of deleterious variants in ROH is not due to inbreeding^50^, but instead to the well-documented ‘Muller’s ratchet’ process, resulting in a lower efficiency of selection in non-recombining genomes.

We also found that some Gene Ontology (GO) categories accumulated higher densities of amino acid-altering variants (Supplementary Note), but the heterogeneous distribution of *N*_*hom*_ and *N*_*alleles*_ across GO categories was virtually identical in RHG, AGR and EUR populations (Supplementary Figs 19 and 20). However, some clinically relevant, disease-related variants, most of them initially discovered in Europeans, were observed at different, measureable frequencies across populations (Supplementary Note and Supplementary Table 10). Together, our analyses indicate that, despite marked differences in the demographic regimes of the African populations examined, these populations have similar mutational burdens, regardless of the average dominance coefficient and the genomic location of the deleterious mutations.

## Discussion

Our study sheds new light on the demographic parameters characterizing the history of African rainforest hunter-gatherers and farmers, and shows that their contrasting demographic histories have not led to major differences in the burden of deleterious variants. Over the years, a number of demographic models have been proposed for these populations, generating conflicting results^34-39,51^. Several studies have dated the divergence of the ancestors of RHG and AGR to ∼60 kya^34-37^, but a recent report, based on 16 whole-genome sequences, inferred a more ancient divergence, 90 to 155 kya, but had only limited power for the estimation of recent population size changes and gene flow due to the small sample size^38^. Our results suggest that RHG and AGR populations diverged between 97 and 140 kya, consistent with an ancient divergence between these human groups^38^. The inclusion of Europeans in our analyses made it possible to test different models for the chronology of major population splits. The similar likelihoods obtained with all models suggest that the ancestors of the RHG, AGR and EUR populations diverged from each other at around the same time (∼85-140 kya), during a period of major climate change (*e.g.*, megadroughts dated between 75 and 135 kya)^52^ that probably promoted population isolation and ancient structure on the African continent^38,51^.

Several other important conclusions about the recent demographic history of African populations can be drawn from our analyses, owing to the combined use of sequencing data from large sample sizes and refined demographic modeling. Regardless of the branching model considered, our inferences indicate that the effective population size of RHG was at least as large as that of the ancestors of AGR for most of their evolutionary past. RHG groups were, therefore, probably more demographically successful, or more interconnected by gene flow, in the past than in more recent millennia, as has also been suggested for the Khoe-San hunter-gatherers of southern Africa^53^. More recently, RHG and AGR populations have undergone very different demographic events, with RHG populations experiencing major size reductions and AGR populations mild expansions, accompanied by a marked increase in gene flow between them. Together, our results support the notion that the traditional mode of subsistence of human populations is correlated with differences in demographic success^31,34,35,39,54^. They extend previous findings by providing the precise parameters characterizing the history of African RHG and AGR over the last 150,000 years.

The impact of population growth and decline on the efficiency of purifying selection and mutational burden in humans has been the subject of intense research over the last few years^8,14-22^. We directly inferred parameters for selection efficiency from the distribution of fitness effects of new nonsynonymous mutations, rather than by simulation-based approximation^15,24^. Our results show that Europeans experienced, on average, weaker purifying selection than Africans, but that the proportions of mutations assigned to different classes of fitness effects did not differ significantly between the African RHG and AGR populations, in either the western or eastern groups. Thus, despite the strong collapse of RHG populations, our findings suggest that the recent nature of this demographic event, together with the historically large *N*_e_ of these populations, has resulted in a selection efficiency similar to that estimated for the expanding farmers.

We also compared the number of deleterious alleles per individual across populations – a statistic that has been shown to be monotonically related to the additive mutation load – accounting for evolutionary sampling variance by resampling sites^27,28^. We found negligible differences in the per-individual number of deleterious alleles between African and European populations. Furthermore, our neutral simulations, which included a strong out-of-Africa bottleneck and a recent expansion for Europeans, reproduced the empirical distribution of synonymous alleles per individual across populations. This result validates our demographic inferences based on independent aspects of the data (*i.e.*, the SFS), and suggests that changes in effective population size alone can account for the observed relative counts of alleles across populations. Additionally, the confidence intervals of ratios of per-individual genotype counts obtained from simulations matched those obtained by resampling sites (Supplementary Table 9), indicating that experimental variation due to factors, such as sequencing coverage, had not inflated the uncertainty of our estimates. Together, these results are consistent with theoretical predictions that the demographic changes experienced by African and European populations are too recent to have had an impact on the mutational load of these populations under an additive model of dominance^23^.

In a scenario in which deleterious variants are partially recessive, homozygous sites would be expected to make the major contribution to mutation load. The number of homozygous functional sites in Europeans was significantly larger than that in Africans, consistent with previous findings^26^, but this excess would not necessarily result in a higher deleterious load in Europeans^28^. Our new empirical data show that African RHG and AGR populations differ only slightly in terms of the number of homozygous functional sites, suggesting equivalent mutational loads in these populations, regardless of the dominance model assumed. Our study revealed differences in the demographic regimes of African populations with different subsistence strategies, but these changes in population size occurred too recently for a difference in mutational burden between populations to have become evident. Furthermore, the strong gene flow inferred between RHG and AGR populations in recent millennia may have attenuated the effect of the strong bottleneck experienced by hunter-gatherers on the efficiency of purifying selection and mutation load.

Extensive exchanges of migrants between human populations have often occurred^55^, and future studies on the dynamics of mutation load in admixed populations should improve our understanding of the impact of gene flow on the distribution of deleterious variants, and, ultimately, the genetic architecture of human diseases.

## Methods

### Population and Individual Selection

In total, we included 317 individuals from the AGR and RHG populations of western and eastern central Africa, and 101 individuals of European ancestry^42^ (Supplementary Note and Supplementary Table 1). Informed consent was obtained from all participants in this study, which was overseen by the institutional review board of Institut Pasteur, France (2011-54/IRB/6), the *Comité National d’Ethique du Gabon*, Gabon (No. 0016/2016/SG/CNE), the University of Chicago, United States (IRB 16986A), and Makerere University, Uganda (IRB 2009-137).

### Exome Sequencing and Quality Controls

We sequenced the exome of 314 African samples, and processed these data together with 101 European individuals^42^. All samples were sequenced with the Nextera Rapid Capture Expanded Exome kit, which delivers 62 Mb of genomic content per individual, including exons, untranslated regions (UTRs), and microRNAs. Using the GATK Best Practice recommendations^56^, read-pairs were first mapped onto the human reference genome (GRCh37) with BWA v.0.7.7 (ref.^57^), and reads duplicating the start position of another read were marked as duplicates with Picard Tools v.1.94 (http://picard.sourceforge.net/). We used GATK v.3.5 (ref.^58^) for base quality score recalibration (“BaseRecalibrator”), insertion/deletion (indel) realignment (“IndelRealigner”), and SNP and indel discovery for each sample (“Haplotype Caller”). Individual variant files were combined with “GenotypeGVCFs” and filtered with “VariantQualityScoreRecalibration”.

As a criterion to remove low quality samples, we required at least 40x of mean depth of coverage (3 excluded samples), 85% of the positions in the BAM file to be covered at 5x minimum (8 excluded samples) and a missingness lower than 5% (1 excluded sample) (Supplementary Fig. 6). In addition, we checked for unexpectedly high or low heterozygosity values suggesting high levels of inbreeding or DNA contamination, and excluded 3 additional individuals presenting heterozygosity levels 4 SD higher than their population average (Supplementary Fig. 6). We thus retained exome data from 400 individuals, with an average depth of coverage of 68x, ranging from 40x to 168x, and an individual breadth of coverage above 5x for 93% of the exome target on average, ranging from 85% to 97%. Finally, we removed indels and discarded from the 768,143 SNPs obtained those that: (i) were not biallelic, (ii) presented missingness above 5% (iii) were monomorphic in our sample, (iv) were located on the sex chromosomes, (v) presented a Hardy-Weinberg test *P*-value < 10^−3^ in each of the populations, and (vi) had an unknown ancestral state using the 6-EPO multi-alignment from Ensembl Compara v59. The application of these quality-control filters resulted in a final dataset of 488,653 SNPs (406,270 SNPs segregating in African populations and 82,383 European-specific SNPs). We intersected the variants with dbSNP build 149 and identified 79,191 previously unreported SNPs.

We also examined whether variation in individual missingness affected the per-individual number of homozygotes (*N*_*hom*_) or alleles (*N*_*alleles*_=*N*_*het*_+2×*N*_*hom*_), and found a significant correlation between individual missingness and these parameters. We thus allowed no missingness and required a depth of coverage of at least 5x for each variant across all individuals, leading to a filtered dataset of 382,786 SNPs. To test the influence of depth of coverage on the detection of SNPs in this new dataset, we explored the correlation between the per-individual mean and variance of coverage and the number of SNPs identified per individual, and observed no significant correlation (Supplementary Fig. 11). This filtered dataset was then employed for all analyses that used per individual genotypes and allele counts.

### Functional Annotation of Variants

To annotate synonymous and nonsynonymous variants in our dataset, we first obtained a bed file with genomic coordinates of exons for each canonical transcript from the UCSC genome browser (http://genome.ucsc.edu/) with the “Table” browser feature track from “KnownCanonical” UCSC genes table. We then used a custom script to parse the concatenated exons of each transcript, codon by codon, and annotate sites within each codon as 0-fold or 4-fold degenerate according to the genetic code. The annotation of Loss-of-function (LOF) variants was performed by Ensembl Variant Effect Predictor (VEP version 84). Only stop-gained and splice disrupting variants were considered. Frameshift variants were not detected because of preliminary filters removing indels. We applied the Loss-of-Function Transcript Effect Estimator plugin (LOFTEE, available at https://github.com/konradjk/loftee) to attribute high (HC) or low (LC) confidence to LOF variants. With LOFTEE, we filtered out LOF mutations known as being false positives, such as variants near the end of transcripts and in non-canonical splice sites, and kept only functional annotations based on the transcript having the highest confidence label.

Additionally, to assess the biological impact of each variant, we used the conservation-based scores computed by the algorithm GERP on an alignment of 35 mammals^59^. Positive GERP Rejected Substitution (RS) scores indicate high degree of conservation and therefore, a high probability for variants at conserved sites to be deleterious. We examined only variants with GERP RS scores greater than 4 because this class has been reported to be under substantial selective constraint^17^ and contained a sufficiently large number of sites in our exome data to have power to detect potential differences between populations.

### Genetic Diversity of Hunter-Gathering and Agriculturalist Populations

We computed several summary statistics on synonymous sites. The Watterson expected nucleotide diversity (*θ*_*W*_), the pairwise nucleotide diversity (*θ*_*π*_), and Tajima’s *D* were calculated based on the site frequency spectrum, with custom R scripts. Confidence intervals and *P*-values for comparisons of these summary statistics between populations were computed by bootstrapping 1000 times and re-sampling sites with replacement.

### Site Frequency Spectra

Our demographic and selection inferences were based on fitting models to folded site frequency spectrum (SFS) data. We used folded SFS in order to avoid ancestral allele misspecification. To obtain the SFS for nonsynonymous and synonymous site classes, we created a VCF file containing variant and invariant sites, kept loci with at least 5x coverage and allowed no missingness across individuals. We then used PGDspider^60^ to transform our VCF files to arp files, and used Arlequin v.2.5 (ref.^61^) to calculate 1D SFS per population, and pairwise 2D SFS.

### Demographic inference

To estimate parameters of demographic models, we used the program *fastsimcoal2* ver. 2.5.2.21 (ref.^43^) (http://cmpg.unibe.ch/software/fastsimcoal2/). *fastsimcoal2* performs coalescent simulations to approximate the likelihood of the data given a certain demographic model and specific parameter values. Maximization of the likelihood is achieved through several cycles of an expectation maximization (EM) algorithm. Therefore, it is critical to perform several simulations to approximate with high precision the likelihood, enough cycles of the EM algorithm to ensure the maximum was reached, and several independent replicate estimations to ensure the global maximum likelihood was found. For all our point estimates, we performed 500,000 simulations, 30 cycles of the EM, and 100 replicate runs from different random starting values. We recorded the maximum likelihood parameter estimates that were obtained across replicate runs. For calculation of confidence intervals (CIs), we fitted the demographic models to resampled SFS data obtained by bootstrapping 100 times by site using Arlequin61. Parameter inference for each bootstrap replicate was achieved by performing 500,000 simulations, 20 cycles of the EM, and 20 replicate runs from different starting values.

We used non-CpG 4-fold degenerate synonymous sites from 21,782 genes for our demographic inferences, having in total 2,383,014 invariant and 223,356 variant sites that passed quality filters. We used these data to generate 1-dimensional (1D) folded SFS for each population, and 2-dimensional (2D) folded SFS for each pair of populations with Arlequin^61^. We fitted two types of demographic models to the SFS data, assuming a mutation rate of 1.36×10^−8^ per-site per generation (*i.e.,* the mutation rate inferred for non-CpG sites)^62^, and a generation time of 29 years^63^. Firstly, three-epoch models of demographic change were fitted to the 1D SFS of each population to obtain rough approximations of the size changes that each population has experienced. Secondly, we fitted complex models including splits, gene flow and size changes to the pairwise SFSs of all the sampled populations. For these computations, we assumed that pairwise SFSs are independent and, therefore, *fastsimcoal2* computed a composite likelihood. Moreover, the maximum likelihood method employed by *fastsimcoal2* assumes that sites are unlinked, therefore it calculates the full likelihood of the data by multiplying the likelihood for each site. Since we used only a few SNPs per gene and, assuming that on average there is no substantial linkage between genes, we expect that the assumption of *fastsimcoal2* that sites are unlinked is reasonable for our dataset.

We used synonymous sites for demographic inference, because they are, among all sites in our exome data, the least likely site class to be under natural selection. We expect, however, that linked selection *(i.e.*, background or positive) might have impacted the diversity and the shape of the SFS and, particularly, the variance of these statistics among sites^64,65^. By using the average SFS across sites for demographic inference, and not variance statistics, we reduced the potential impact of linked selection on our inference. Moreover, the effects of background selection on introducing bias for inference of population size changes are most severe when the strength of linked selection is moderate (*N*_*e*_*s* =2-20)^64^. Our inferred DFEs show that the proportion of nonsynonymous mutations in this *N*_*e*_*s* range is rather small (10-20 %), thus we would not expect a substantial bias on the population size change estimates. For migration rates, strong background selection can actually improve estimation^64^. Furthermore, in humans, who have a large recombining genome, we would expect positive selection to impact demographic inference only if it is highly pervasive^65^, a scenario that is highly unrealistic^66^. Finally, we expect that the confidence intervals on the demographic parameters, which were computed by bootstrapping by site, should reflect some of the uncertainty over the parameter estimates introduced by background selection or selective sweeps.

### Distribution of Fitness Effects of New Mutations (DFE)

We used methods implemented in *DFE-α* v.2.15 (ref.^46^) and *DoFE* (ref.^48^) to infer the DFE of new nonsynonymous mutations. *DFE-α* fits a demographic model to a class of sites that is assumed to be neutral and, conditional to the demographic model inferred, fits a gamma DFE model to the SFS of the focal class of sites. *DoFE* does not assume a demographic model and instead fits nuisance parameters to the neutral SFS and a gamma DFE model to the focal SFS. We used the synonymous and nonsynonymous SFS as neutral and focal classes, respectively. Both methods infer the mean and the shape of the gamma distribution, and we also interpolated the proportion of mutations that are assigned to four *N*_*e*_*s* ranges (0-1, 1-10, 10-100, >100) corresponding to neutral, weakly selected, strongly selected and lethal mutations. We calculated confidence intervals for *DFE-α* by bootstrapping by site 100 times, and for *DoFE*, we calculated the 0.025 and 0.975 quantiles of the posterior distribution of the estimated parameters.

### Estimating Mutation Load

Mutation load was approximated using the number of putatively deleterious mutations per individual in several functional categories: nonsynonymous variants, variants with GERP RS greater than 4 (ref.^59^), and LOF mutations. We counted the number of deleterious alleles in each individual as *N*_*alleles*_=*N*_*het*_+2×*N*_*hom*_, with *N*_*het*_ and *N*_*hom*_ corresponding to the numbers of heterozygous and homozygous genotypes, respectively^17,23^. We also performed 100 coalescent simulations, with *fastsimcoal2* (ref.^43^), to predict the expected neutral variation per individual of the five studied populations, across the three branching models. To do so, we computed on the simulated data the number, per individual, of heterozygous genotypes *N*_*het*_, homozygous genotypes *N*_*hom*_, and alleles *N*_*alleles*_, using the files produced by *fastsimcoal2*, containing the maximum likelihood point estimates for the parameters of each demographic model. The significance of population differences in per-individual observed and simulated genotype counts was assessed by estimating the confidence interval of ratios between population pairs. Confidence intervals were calculated by paired bootstrapping: we split the SNP data into 1000 blocks and resampled, with replacement, 1000 times. This approach takes into account the variance introduced by demographic processes^27,28^.

### Runs of Homozygosity (ROH)

We searched for ROH along the genome of all individuals using the sliding window approach implemented in PLINK^67^ on the SNP genotyping data. The whole genome of each sample was explored using sliding windows of 50 SNPs, and ROH were detected if the 50 SNPs were homozygous with the possible exception of two heterozygous and five missing genotypes. Various minimum lengths were tested to define ROH regions. Finally, we allowed any length of ROH regions and discerned inbreeding from strong linkage disequilibrium (LD) on homozygous segments by classifying ROH in two size classes, adapted from Pemberton *et al*. (2012)^68^. Class A ROH were 0-0.5 Mb in length, and are attributable to high levels of local LD. Class B ROH were > 0.5 Mb in length, resulting from background relatedness due to limited population size or inbreeding.

### Hotspots and Coldspots of Recombination

We defined regions of high recombination rates (HRR) and coldspots of recombination (CS) using the full list of autosomal regions provided in ref.^69^. We obtained the intersections of the positions of each SNP in our dataset with the coordinates of HRR and CS regions, and assigned them to one of the two categories.

### Gene Ontology Categories

We assigned Gene Ontology (GO) categories for a set of 16,290 genes from the 21,782 genes found in the exome target using the GOSeq package^70^. From the 18,585 GO categories represented, we selected 16 functions including at least 200 genes, and being biologically relevant in the context of comparisons between hunter-gathering and farming populations. We reported 1,371 genes in “Locomotion”, 1192 genes in “Immune response”, 1,142 genes in “Lipid metabolic process”, 928 genes in “Embryo development”, 850 genes in “Reproduction”, 814 genes in “Cardiovascular system development”, 766 in “Sensory perception”, 766 in “Response to hormone”, 591 in “Brain development”, 576 in “Protein catabolic process”, 490 in “Muscle structure development”, 323 in “Skin development”, 300 in “Response to oxidative stress”, 285 in “Developmental growth”, 217 in “Aging”, 208 in “Glucose metabolic process”. In order to compare mutation load results across categories, we divided all scores per the length of the coding sequence of all genes included in each category.

### Clinically-relevant Variants

In order to detect clinical variants with validated pathogenicity in the exomes of African populations, we intersected the set of 406,270 SNPs segregating in RHG and AGR groups with the curated ClinVar database^71^ available here: https://www.ncbi.nlm.nih.gov/clinvar/docs/maintenance_use/. We only considered ClinVar entries with the most supported evidence for clinical significance, recorded as “5 - Pathogenic”. We therefore identified a total of 334 pathogenic variants distributed in 251 genes, and mostly segregating at a derived allele frequency lower than 3%.

## Data availability

The newly generated exome sequencing data for the Central African rainforest hunter-gatherers and agriculturalists (*N*=300) have been deposited in the European Genome-phenome Archive under accession code EGASXXXXXXX. Exome sequencing data for the European population (*N*=100) are available under accession code EGAS00001001895.

## Acknowledgments

We thank all of the participants for providing the DNA samples used in this study. We thank the Paleogenomics and Molecular Genetics Platform of the MNHN-Musée de l’Homme for technical assistance in DNA sample preparation. We thank Guillaume Laval, Laurent Excoffier and Maxime Rotival for helpful discussions, and Nicolas Joly for help in computational resources. This work was supported by the Institut Pasteur, the *Centre National de la Recherche Scientifique* (CNRS), and the *Agence Nationale de la Recherche* (ANR) Grant “AGRHUM” (ANR-14-CE02-0003-01).

### Author contributions

M.L. and A.K. designed the analytical approach and performed the analyses, with input from E.P. and L.Q.-M. H.Q. and C.H. performed experiments, P.M.-D, J.-M.H., A.F., G.H.P., L.B.B. and P.V. collected the samples. E.P. performed analyses. L.Q.-M. conceived the study, with input from E.P., and obtained funding. The manuscript was written by M.L., A.K., E.P. and L.Q.M, with input from all authors.

### Competing financial interests

The authors declare no competing financial interests.

